## Supplementary material for "Jagged-mediated lateral induction patterns Notch3 signaling within adult neural stem cell populations": Full supplementary material

**Ortica et al.**

### Supplementary Figures and Supplementary Figure Legends

**Figure S1. Details on the BAC and Crispr recombination procedures. A.** Left: schema of the Notch3 protein with some regions of interest: Epidermal Growth Factor (EGF) repeats, Lin-12/Notch repeat (LNR) and Heterodimerization (HD) domain on the extracellular domain, Nuclear Localization Signals (NLS), Ankyrin repeats (ANK) domain and two Proline/glutamic acid/serine/threonine rich domains (PEST1/2) in the intracellular domain. Right: scheme of the Notch3 fusion proteins expected after transgenesis: a green fluorescent protein (GFP or Azami Green -AG-) followed by a ribosome skipping P2A domain and a nuclear RFP (nlsRFP, magenta) are inserted at the C-terminus, partially or totally disrupting Notch3 PEST domains. **B.** Aminoacidic sequence of the zebrafish Notch3 intracellular domain (N3ICD). Orange*:* putative gamma-secretase cleavage site; blue: ANK domain; magenta*:* PEST domains; green*:* sites of insertion of the fluorescent reporter proteins for the TgBAC$^{n3\text{-}GFP/+}$ and the TgKI$^{n3AG/+}$ lines. **C.** Scheme of modifications performed on BAC CH211-69E21 carrying the zebrafish *notch3* promoter and gene sequence, used to generate the TgBAC$^{n3\text{-}GFP/+}$ line. First, the *pTARBAC2.1* vector was modified at the *loxP511* site to insert a CFP under the beta-crystallin promoter as a transgenesis marker, as well as the iTol2-Amp cassette for random integration in zebrafish genome and Ampicillin selection. Then, a cassette containing the GFP-P2A-nlsRFP sequence was inserted in frame into *notch3* exon33. Removal of the Zeocine Resistance (ZeoR) cassette used for selection could not be performed. **D.** Top: guideRNA (gRNA) sequences tested to generate the TgKI$^{n3AG/+}$ line; and their scores as noted on the CRISPOR web tool [1] (specificity score, predicted efficiency according to Moreno-Mateos parameters [2], number of off-targets for 0-1-2-3-4 mismatches and number of off-targets followed by a PAM for 0-1-2-3-4 mismatches). Bottom: position of gRNA2 and gRNA4 (in green) relative to PEST sequences (magenta) and STOP codon (red) of the *notch3* gene (start and end positions relative to [NCBI Reference Sequence: NC_007114.7] are indicated). **E.** A T7 Endonuclease1 test was performed on PCRs obtained from pools of 20 embryos injected with gRNA2, gRNA4 or not injected. Endonuclease1 cleaves only not-perfectly matched DNA, as it results from CRISPR/Cas9-generated indels. Both gRNA induce indels, but gRNA4 shows a higher percentage of cut (two fragments – black arrows) over uncut (blue arrow) DNA (estimated to 50%), and was selected to generate the TgKI$^{n3AG/+}$ line. **F.** Two rounds of injections in *casper* fish were performed using gRNA4, Cas9 protein and donor vector as in panel B. Percentages of knock-in efficiency and transmission to the first generation are indicated.

**Figure S2. Notch3 fusion alleles report the different subcellular localizations of the Notch3 protein or protein fragments and their intracellular trafficking. A.** Chromogenic in situ hybridization (ISH) on 48 hours post fertilization (hpf) larvae show that the expression patterns of *GFP* and *AzamiGreen* are similar to *notch3* in wild-type larvae. Scale bars: 200 µm. **B**. Left: schematic of the Light sheet microscope imaging angle of a 30hpf TgBAC$^{n3\text{-}GFP/+}$ embryo. Right: snapshot of an optical cross section across the embryonic forebrain, with Notch3-GFP in green and nlsRFP in magenta (V = Ventricle, A-P = Anterior-Posterior axis). Scale bar: 20 µm. The white arrow indicates Notch3-GFP at the membrane and the blue arrow indicates GFP nuclear signal. **C.** Snapshots of a short confocal movie of a 30hpf TgBAC$^{n3\text{-}GFP/+}$ living embryo (Notch3-AG in green). The first image illustrates the whole forebrain region that was recorded, and following images are magnifications of the indicated area (white square) at different times points (time in seconds indicated at the bottom of each image). V = Ventricle, A-P = Anterior-Posterior axis. Scale bar: 20 µm. Yellow and orange arrows show two moving green dots. **D,E.** Whole-mount dorsal confocal views of the adult pallial ventricular zone in each transgenic line, immunostained for endocytic vesicles markers: early endosomes: Rab5, late endosomes: Rab7, recycling endosomes: Rab11, and lysosomes: Lamp1. For each image, a detail of an orthogonal section at higher magnification is shown in inset. White arrows indicate colocalization of Notch3-fusion proteins (green) and endocytic vesicle markers (magenta) (Zo1 in cyan). The apparently different distribution behavior of Notch-GFP (absent in sorting endosomes) and Notch3-AG remains to be interpreted. GFP is reversibly quenched at acidic pH, which has led to differential tracking of GFP versus mCherry fluorophores in Drosophila [3,4]. IHC will be sensitive if quenching is associated with a denatured protein structure. More resolutive imaging would also be needed. Scale bars: 5 µm.

<u>Figure S3.</u> **Functionality of Notch3 fusion alleles in embryos and larvae. A,B.** A 20-minute BrdU pulse was applied to WT, $TgBAC^{n3\text{-}GFP/+}$ and $TgKI^{n3\text{-}AG/n3\text{-}AG}$ embryos at 30hpf. **A.** Forebrain cross sections (10 µm-thick) immunostained for BrdU (magenta) and counterstained with DAPI (cyan) (V = ventricle, A-P = Anterior-Posterior axis). Scale 20 µm. **B**. Proliferation rates (percentage of $BrdU^{pos}$ cells in the area boxed in white in **A**) between genotypes. **C-E.** Quantification of neuronal production in WT, $TgBAC^{n3\text{-}GFP/+}$ and $TgKI^{n3\text{-}AG/n3\text{-}AG}$ embryos at 29 and 48hpf. **C**. Forebrain cross sections (10 µm-thick) at 29hpf immunostained for HuC (white) and counterstained with DAPI (cyan) (V = ventricle, A-P = Anterior-Posterior axis). Scale 20 µm. The same region was compared between brains (white square) to evaluate neuronal production at 29hpf and 48hpf. **D,E**. Neuronal production (percentage of $HuC^{pos}$ cells among all cells in the area boxed in white in (C) -or an equivalent area at 48hpf-) between genotypes (Welch's ANOVA test, p-value=0.48 at 19hpf and 0.16 at 48hpf, ns: non-significant). **F-H.** Quantification of radial glia (RG) activation in WT, $TgBAC^{n3\text{-}GFP/+}$ and $TgKI^{n3\text{-}AG/n3\text{-}AG}$ larval pallia at 8dpf. **F**. Images: left: example of a 50 µm-thick vibratome section across the telencephalon immunostained for Blbp (white) and Pcna (yellow) and counterstained with DAPI; middle and right: individual fluorescence channels. RG: $Blbp^{pos}$ (white); neural progenitors (NP): $Blbp^{neg}$; proliferating cell: $Pcna^{pos}$ (yellow). Scheme: schematic lineage progression depicting quiescent (q) and activated (a) RG, NPs, and neurons (N). Activated cells depicted with a yellow nucleus. Scale bar: 20 µm. **G,H.** Percentages of activated RG ($Blbp^{pos}$;$Pcna^{pos}$ among $Blbp^{pos}$ cells) compared between WT and $TgKI^{n3\text{-}AG/n3\text{-}AG}$ (**G**) (Wilcoxon text, p-value=0.8, ns), or WT and $TgBAC^{n3\text{-}GFP/+}$ (**H**) (Wilcoxon test, p-value=0.1066). **I.** Morphological phenotype of 7dpf larvae from a cross between a $TgBAC^{n3\text{-}GFP/+}$; $n3^{fh332/+}$ fish and a $n3^{fh332/+}$ fish. $GFP^{pos}$ larvae ("green") appear normal, while a measurable proportion of $GFP^{neg}$ larvae ("non-green") display a curved phenotype at 7 dpf, similar to $n3^{fh332/fh332}$ larvae. **J.** Percentage of surviving larvae resulting from a cross between a $TgBAC^{n3\text{-}GFP/+}$ $n3^{fh332/+}$ fish with a $n3^{fh332/+}$ fish. $n3^{fh332/fh332}$ larvae die between 6dpf and 10dpf (grey line). The presence of the fluorescent $TgBAC^{n3\text{-}GFP/+}$ copy can completely rescue the mortality (blue line).

<u>Figure S4.</u> **Spatial analysis of N3ICD levels in link with apical area size and in $TgBAC^{n3\text{-}GFP}$ fish. A,B.** Scatter plots of N3ICD levels (y axis, normalized to nuclear volume - conc°- or in absolute values -sum-) relative to apical area size (x axis, in $\mu m^2$) in two pallial hemispheres of a $TgKI^{n3AG/+}$ fish (same as Fig.3) (**A**) and one pallial hemisphere if a $TgBAC^{n3\text{-}GFP/+}$ fish (**B**). Each dot represents a cell, and background colors (purple, green, yellow) are the same color-coded cutoffs as in Fig.3. Red shades inside the dots measure N3ICD levels summed in the immediate neighborhood of each cell. **C.** Superposed neighborhoods of all $N3ICD^{low}$ cells in a $TgBAC^{n3\text{-}GFP/+}$ fish. For each cell in this group, all the cells surrounding it at a maximal radius of 10µm are plotted at their position with respect to the original cell, color-coded as in B. **D.** Deviation from marked L-functions from an uncorrelated model as a function of the distance (radius in micrometers, x axis) from cells of low N3ICD values (scheme on left). Identity of the mark from top to bottom: N3ICD conc° per nucleus, N3ICD sum per nucleus, apical area. n=256 cells from 1 brain (2 hemispheres). Dashed lines indicate the hypothesis rejection threshold. p-values: N3ICD conc°: 0.01, N3ICD sum: 0.08, apical area: 0.85.

<u>Figure S5</u>. **Expression of Notch ligands and interpretation of Notch3 signaling levels in the adult pallial NSC population. A-G.** Expression levels of the different Notch ligand genes, plotted on the

scRNAseq UMAP of neural stem and progenitor cells of the adult pallium in WT fish (left) [5], and revealed by smFISH (green, magenta or yellow) together with IHC for Pcna and Zo1 (white) on whole-mount confocal dorsal views of adult pallia (right). Scale 10 µm. **H.** Top: comparison of *jag1b* (magenta, left) and *jag2b* (yellow, right) expression revealed by smFISH on *Tg(gfap:GFP)* adult pallia together with IHC for GFP (green) and DAPI counterstaining (orthogonal sections of the ventricular zone). Note the apical localization of *jag1b* transcripts (in NSC, $GFAP^{pos}$ cells) and the deeper localization of *jag2b* transcripts (mainly in $GFAP^{neg}$ cells). Scale bar: 20 µm (A-B = apicobasal axis). Bottom: distribution of *jag1b* (left) and *jag2b* (right) expression on scRNAseq UMAP of all cells of a WT adult pallium (with cell types indicated [5]). *jag1b* expression is mainly found in the RG population, while *jag2b* is weakly expressed in RG and rather expressed in neurons. **I.** Interpretative scheme of the variations of nuclear N3ICD levels in NSCs during quiescence and as they progress along the lineage (over time from left to right). NSCs (hexagons) are represented from their apical surface, in scale, with a green plasma membrane (Notch3-fluo). Green shades in cell nuclei (circle in center of hexagons) and graded triangles below reflect nuclear N3ICD levels. $dla^{neg}$ and $dla^{pos}$ NSCs are shown in white and blue, respectively, which progressively higher *dla*:GFP intensities along lineage progression. Cell division events are indicated by black dots, they are separated by phases of quiescence. Lineage progression is drawn from [6], and N3ICD levels are based on Fig.4. Of note, they were analyzed at the population level and not along an individual lineage as depicted here.

**<u>Figure S6.</u> Notch3 signaling in relation with *dla* expression in neighboring cells and with *jag1b* transcription. A,B.** Quantification of nuclear N3ICD-AG levels as in Fig.4J,K, and classification of each cell according to its number of $dla^{pos}$ neighbors (**A** – Spearman's ρ=0.05 ) or its total number of neighbors (**B** – Spearman's ρ=0.36) (Z-score of N3ICD pixel sum; 3 brains, n=174 cells in total). **C.** Scheme of the *jag1b* locus with position of RPBj binding sites within the regulatory upstream sequence. Single sites are indicated in pink, and a head-to-head site known to be efficiently targeted by NICD [7] is indicted in blue. **D.** Scheme depicting the expected effect of a mild LY411575 treatment (lowering γsecretase activity) on the expression of Notch ligands in the case of lateral inhibition (top, increase of ligand's expression) vs lateral induction (bottom, decrease of ligand's expression).

**<u>Figure S7.</u> *hey1* expression is sensitive to Jag1b/Notch3 signaling. A.** Expression of *hey1* revealed by smFISH (yellow) in a whole-mount adult WT adult pallium (dorsal confocal view) together with IHC for Zo1 (white) and Sox2 (cyan). Scale bar: 20 µm. **B.** Corresponding quantification of *hey1* transcripts (number of smFISH dots per cell; 1 brain, n=118 cells). **C.** Expression of the Notch ligand *jag1b* (magenta) and the Notch target *hey1* (yellow) revealed by smFISH in a whole-mount adult WT adult pallium (dorsal confocal view) together with IHC for Zo1 (white). Some cells express only one player (magenta and yellow arrows), others show co-expression of both (white arrow). Scale bar: 10 µm. **D.** Corresponding scatterplot showing the relative distribution in each cell of (C). There is a weak cell-autonomous correlation between ligand and target expression (n=218 cells, 1 brain; Spearman's ρ=0.38). **E.** Scatter plot of N3ICD levels (Z-score of N3ICD pixel sum) and *hey1* transcription (smFISH, number of dots per cell) in the adult pallium of a $TgBAC^{n3\text{-}GFP/+}$ fish. There is a direct cell-autonomous correlation between the two parameters (n=153 cells from 2 brains; Spearman's ρ=0.15). **F.** Boxplot of *hey1* expression levels (smFISH, number of dots per cell) in $Pcna^{neg}$ vs $Pcna^{pos}$ cells in WT adult pallia. *hey1* expression is significantly lower in proliferating cells (Wilcoxon rank sum test, p-value=2.053e-09) (n=336 cells from 2 brains). **G.** Expression of *hey1* (smFISH, yellow), together with IHC for Zo1 (white) and Sox2 (cyan) in the pallial ventricular zone of $TgBAC^{n3\text{-}GFP/+}$ adults treated with 10 mM LY-411575 or solvent only (DMSO) for 24h (dorsal whole-mount confocal views). Scale bar: 20 µm. **H.** Corresponding quantification of *hey1* expression levels (mean of number of dots per cell) in DMSO vs LY conditions (DMSO: n=2 brains, LY: n=3 brains). **I.** Schematic of the experiment assessing Jag1 knockdown effect on *hey1*: TgKI$^{n3\text{-}AG/+}$ fish were injected with *jag1bMO* or *ctrlMO* followed by a three-day chase before *hey1* smFISH detection. **J.** Corresponding quantification of *hey1* expression levels (mean of number of dots per cell) in the two MO conditions. A weak decrease of *hey1* expression after 72h downregulation of *jag1b* translation could be observed (*ctrlMO*: n=321 cells from 2 brains, *jag1bMO*: n=377 cells from 2 brains).

**A**

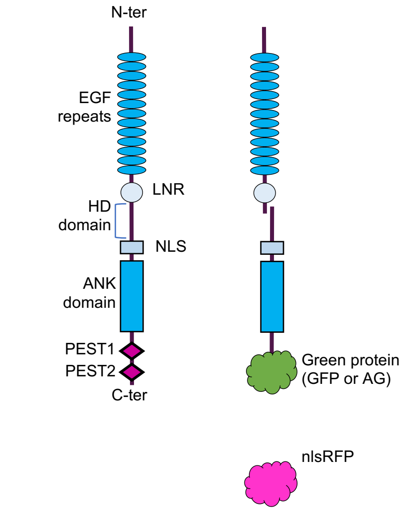


**B**

**N3ICD**

VILMVGMLIARRKREHSTLWFPEGFFLKKETSSNKNRREPVGQDSLGMK
HMPKTVEESLLADHSDQWIDTDCPEAKRLKVEEPSILSDGEDAVDSRQ
WTQHHLAAADIRMPPSMALTPPQGEFDSDCMDVNVRGPDGFTPLMLA
SFCGGGLEPEVTEDDDSDESSANIISDLIYQGASLAAQTDRTGETALHLA
ARYARADAAKRLLDAGADANAQDNTGRSPLHAAVAADAQGVFQILIRNR
ATDLDARMYDGSTALILAARLAVEGMVEELITCHADVNAVDEIGKSALHW
AAAVNNVEATIALLKNGANKDMQDLKEETPLFLAAREGSCEAVKVLLAH
FANREITDHMDRLPRDIAQERMHHDIVQLLDEYNTVRSPQGHAGAGHHL
SGSHTLSPLMCPPSNFLQGLKSTPQGKKNRRPGAKGIGGQHASGLKDS
AKGRNKRLTLDMQSALLESSVTLSPVDSLDSPRGGASNAGYVTNPTSP
AAMPSPGLFHSSMSVPNTPMVHSSILDSTSPFSVSLAQLSDLGDGGLS
MQGRVGMQGGGVNPGPHNYVMNAGQMSLSMGMVSPVSVPFDWHNR
MPPSSQCSQPMISMVHPVSSQAGMLPQTPTLQSSMLLHQQVFRNAQQ
PILQPTPVSSTPSISPVKLPPIAEQQPQQLHSHTLTNPNLSRMATSTPPT
PQQAQPPPAFYQPQQPQAQPPQAPTAPQATQAQAIPSQAPPAQVSGS
TAGPEDYPTPPSQHSYSSAMDATPKHYLHVPNEHPYLTPSPESPEPWS
SPSPHCVSDWSDSTPSPAVTAPSQTQISQVQESNGKMQVFA

**TgKI**$^{n3AG/+}$ **TgBAC**$^{n3-GFP/+}$

**C**

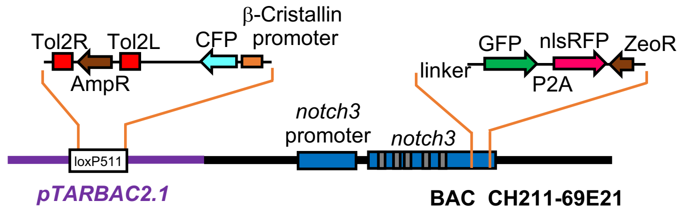


**D**

| | Sequence | Spec. Score | Eff. Score | Off-targets | Off-targets +PAM |
|---|---|---|---|---|---|
| **gRNA2** | GGAGCTGTGACAGCTGGGCTGGG | 94 | 63 | 0-0-0-2-35 | 0-0-0-0-1 |
| **gRNA4** | GGGGTAATCCTCTGGGCCTGCGG | 99 | 77 | 0-0-0-0-3 | 0-0-0-0-0 |

***(28911)***CTGCCTCCCATTGCTGAGCAGCAGCCGCAGCAACTTCACAGCCACACCCTCACCAACCCCAACCTCTCCCGCATGG
CCACCTCCACTCCCCCAACTCCACAGCAGGCACAGCCTCCTCCAGCTTTCTACCAACCACAACAGCCACAAGCTCAGCCGC
CGCAGGCCCCGACAGCCCCTCAGGCAACGCAAGCACAAGCTATCCCATCTCAGGCACCCCCAGCACAGGTGAGCGGCAG
CA**CCGCAGGCCCAGAGGATTACCCC**ACACCACCTTCCCAGCACAGCTACAGCTCAGCCATGGATGCAACGCCTAAGCATTA
TCTGCATGTGCCCAATGAACATCCTTACCTGACCCCTTCTCCAGAGTCTCCAGAGCCCTGGTCCAGCCCTTCCCCTCACTGT
GTGTCTGATTGGTCTGACTCCACA**CCCAGCCCAGCTGTCACAGCTCC**TTCTCAAACCCAGATTTCACAAGTCCAGGAGTCCA
ACGGGAAGATGCAGGTGTTTGCT***TGA***GCAGCCCAACCTACCTTAAAT***(29438)***

**E**

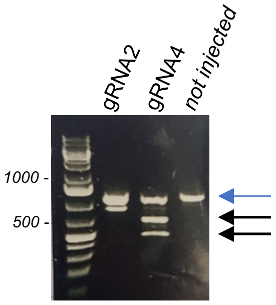


**F**

| | Efficiency | Founders | Transmission rate |
|---|---|---|---|
| **Injection #1** | 3,3% | 1/60 | 3,3% |
| **Injection #2** | 2,8% | 1/36 | 20% |

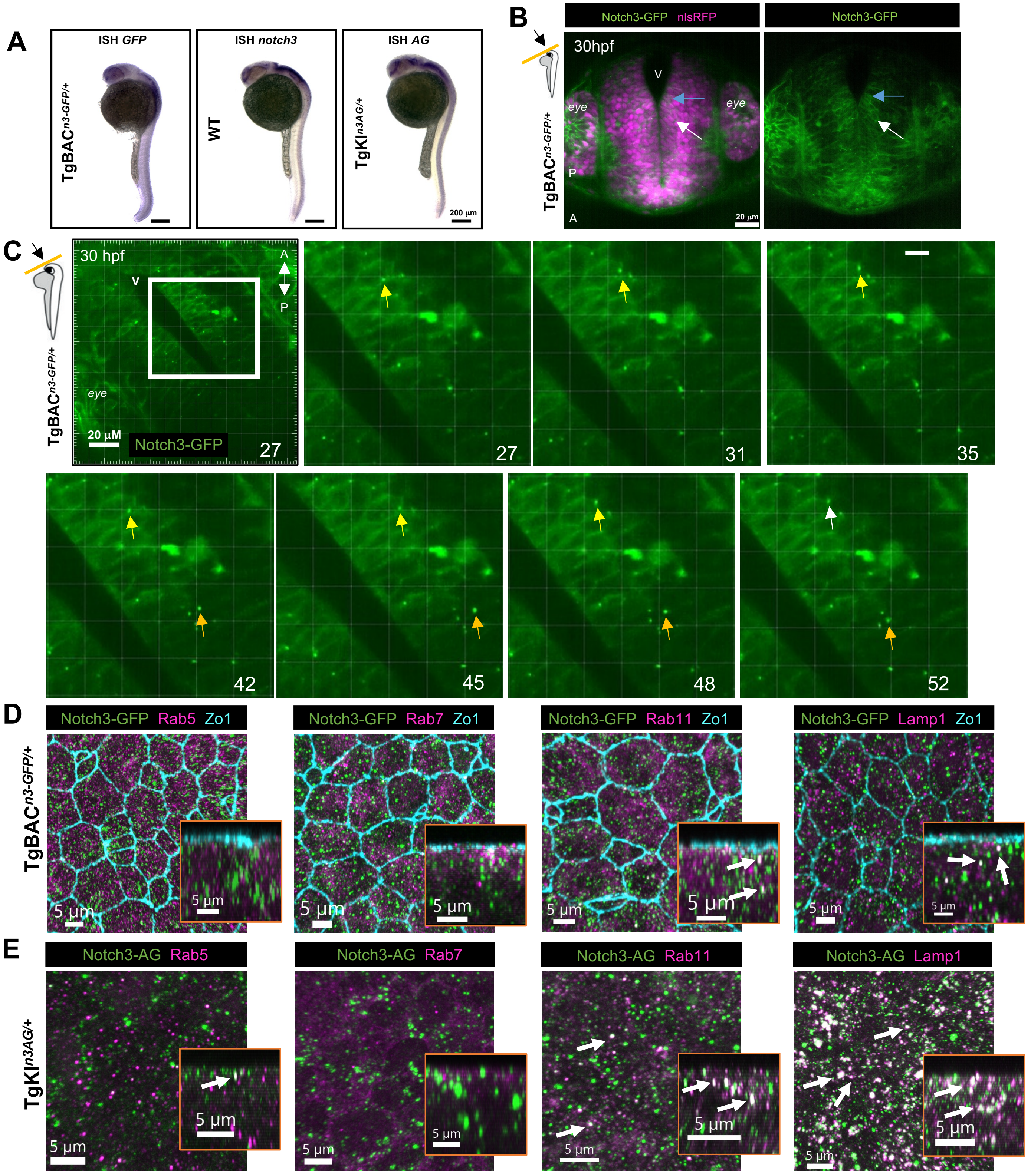


Ortica et al., Figure S2

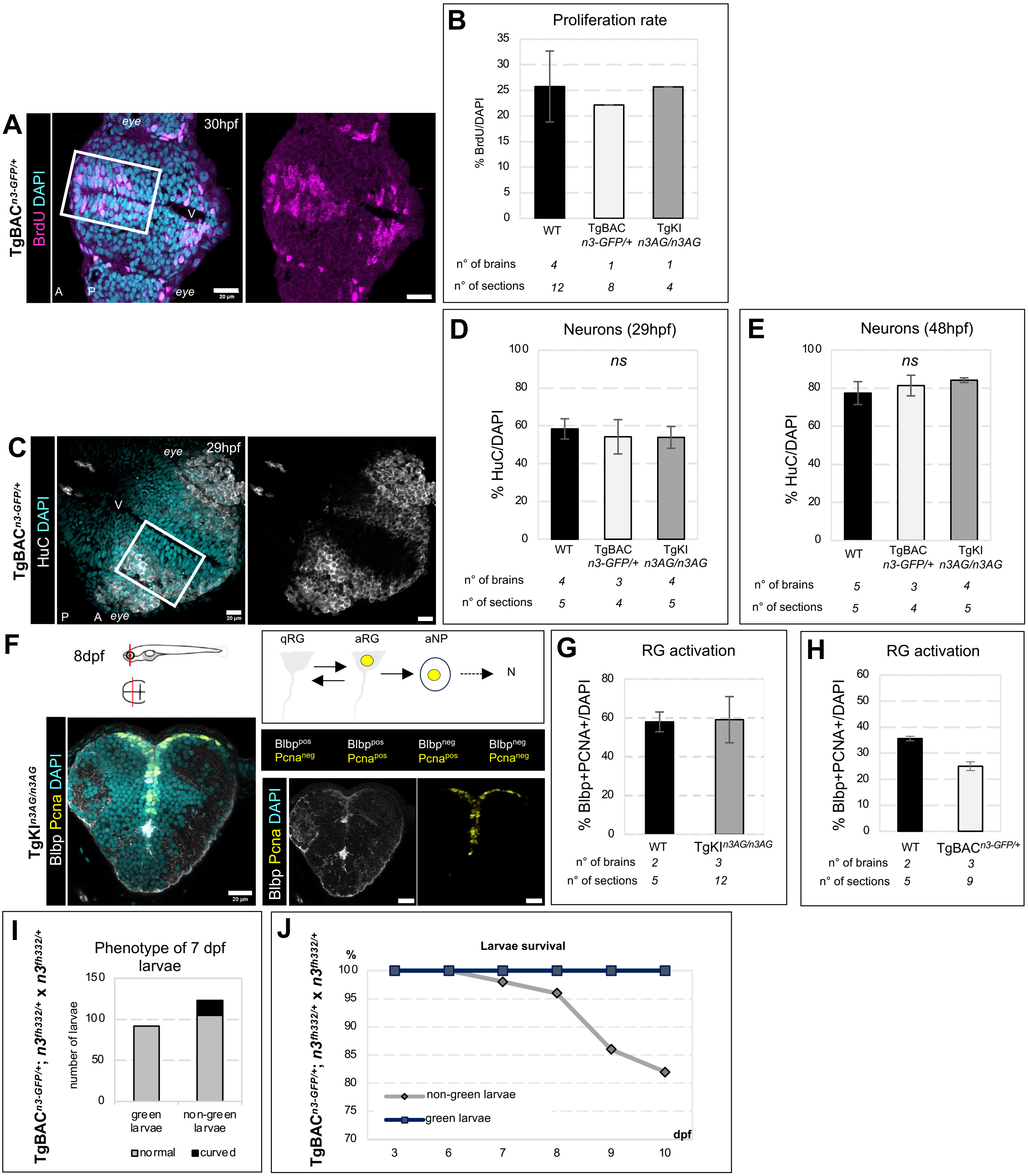


Ortica et al., Figure S3

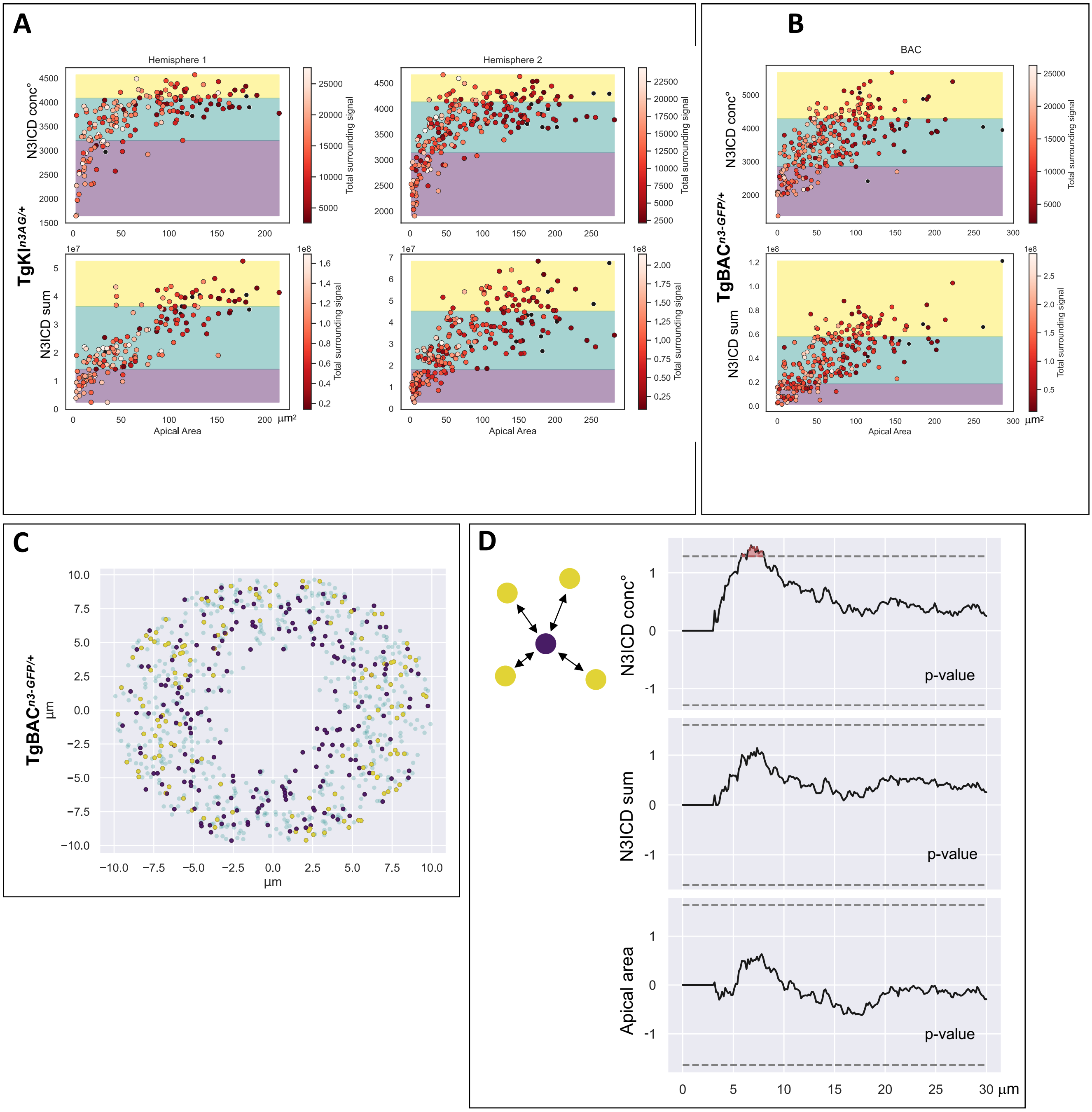
A
Hemisphere 1
Hemisphere 2
N3ICD conc°
N3ICD sum
Total surrounding signal
Apical Area
μm²
TgKI$^{n3AG/+}$
B
BAC
TgBAC$^{n3-GFP/+}$
C
μm
D
N3ICD conc°
N3ICD sum
Apical area
p-value
30 μm

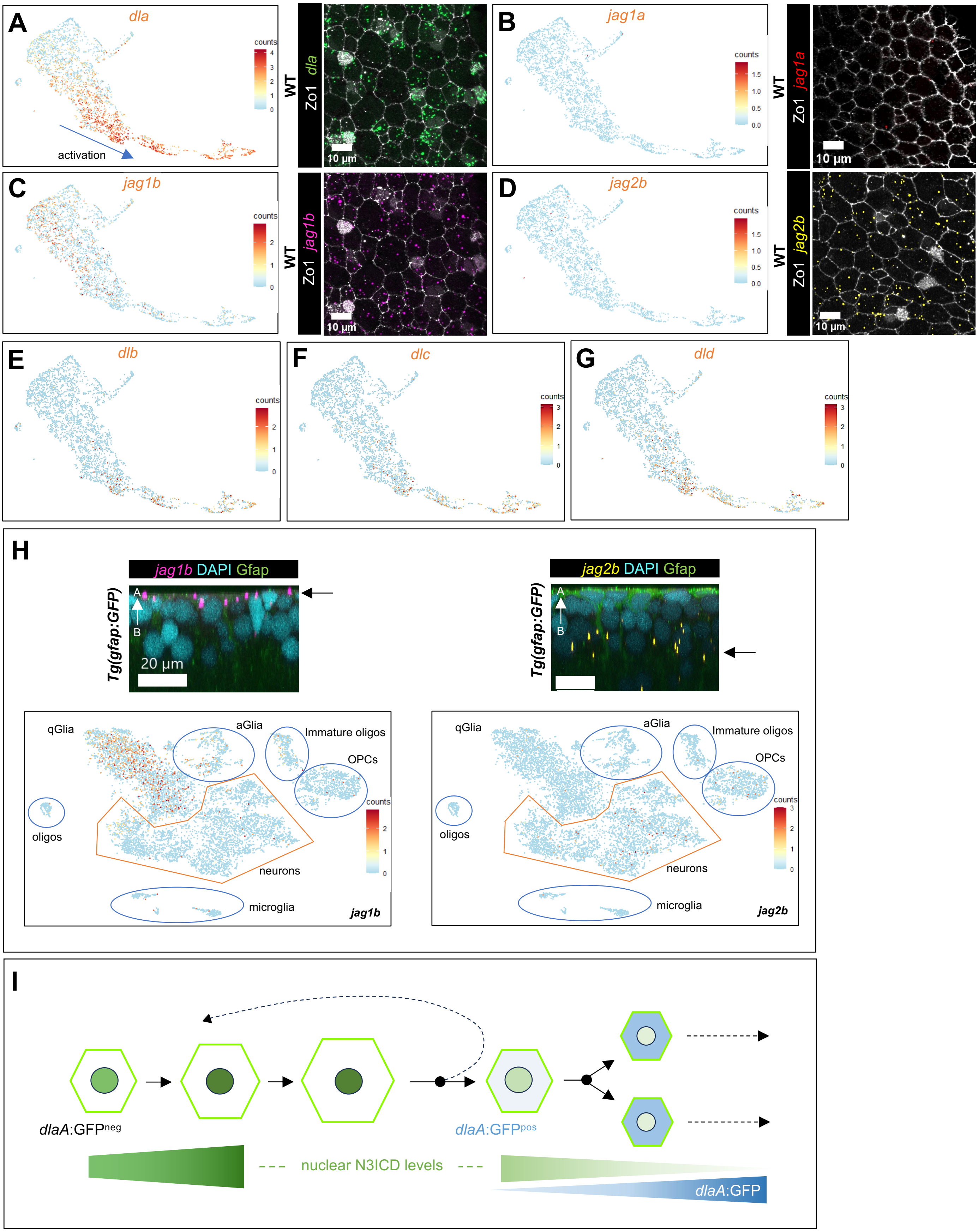


Ortica et al., Figure S5

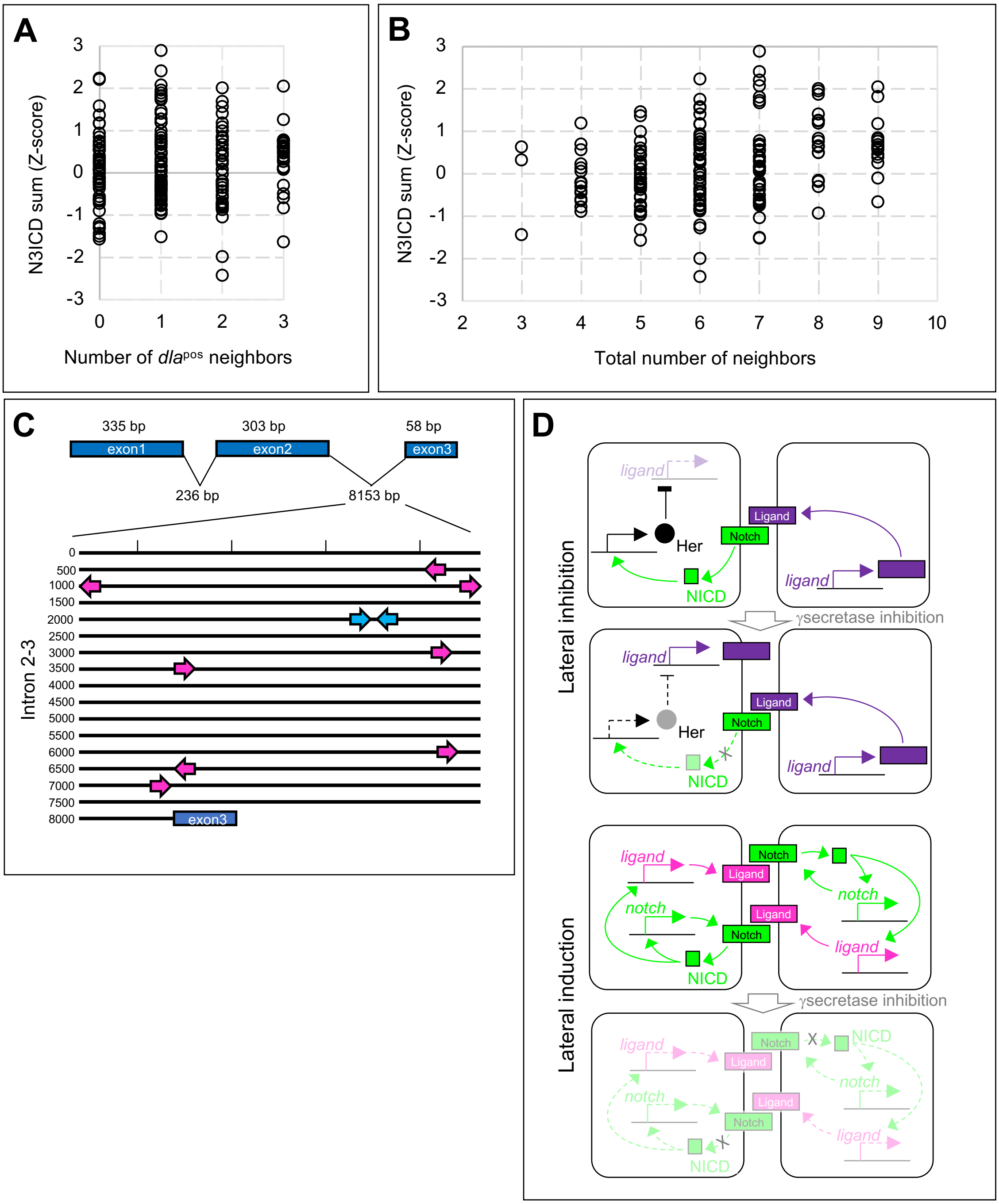


Ortica et al., Figure S6

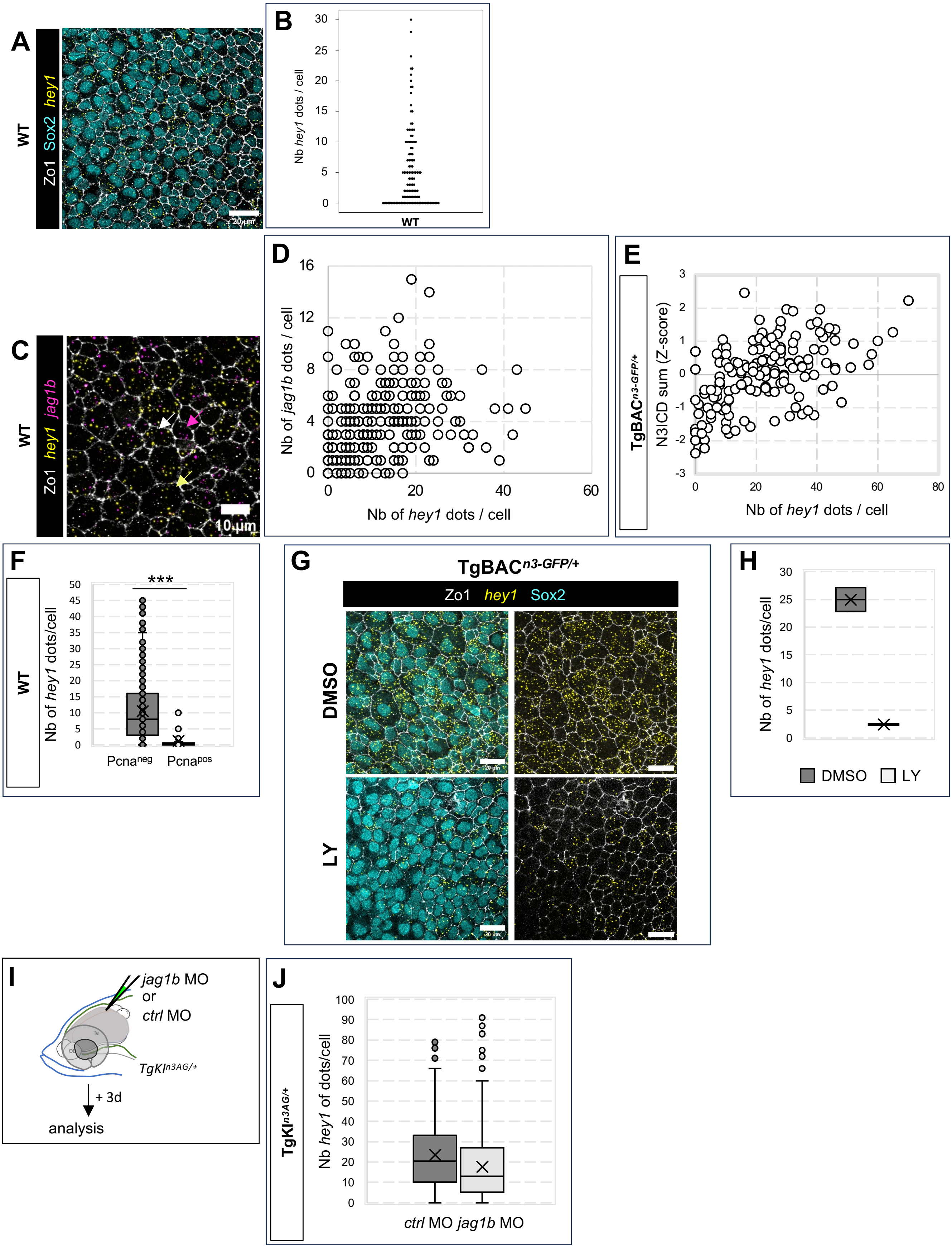


Ortica et al., Figure S7
